# Emergent properties of a mitotic Kif18b-MCAK-EB network

**DOI:** 10.1101/2020.08.09.242602

**Authors:** Toni McHugh, Julie P.I. Welburn

**Affiliations:** Wellcome Trust Centre for Cell Biology, School of Biological Sciences, University of Edinburgh, Edinburgh EH9 3BF, Scotland, UK

**Keywords:** motor, mitosis, microtubule

## Abstract

The precise regulation of microtubule length during mitosis is essential to assemble and position the mitotic spindle and segregate chromosomes. Prior work has identified key molecular players in this process, including the kinesin-18 Kif18b and the kinesin-13 Kif2C/MCAK, which both promote microtubule depolymerization. MCAK acts as a potent microtubule depolymerase diffusing short distances on microtubules, while Kif18b is a mitotic processive motor with weak depolymerase activity. However the individual activities of these factors cannot explain the dramatic increase in microtubule dynamics in mitosis. Using *in vitro* reconstitution and single molecule imaging, we demonstrate that Kif18b, MCAK and the plus-end tracking protein EB3 act in an integrated manner to potently promote microtubule depolymerization. We find Kif18b acts as a microtubule plus end delivery factor for its cargo MCAK, and that Kif18b also promotes EB accumulation to plus ends independently of lattice nucleotide state. Together, our work defines the mechanistic basis for a cooperative Kif18b-EB-MCAK network with emergent properties, that acts to efficiently shorten microtubules in mitosis.

## Introduction

Throughout eukaryotes, length control of microtubule polymers is essential. Microtubule length is controlled through the regulation of microtubule dynamic parameters by motors and microtubule-associated proteins (Mitchison and Kirschner, 1984). At the onset of mitosis, microtubule catastrophe rate increases dramatically, leading to disassembly of the interphase microtubule network (Piehl and Cassimeris, 2003). These dynamic properties of microtubules are essential to enable the assembly of the mitotic spindle, made of shorter antiparallel microtubules, and promote chromosome alignment. In most eukaryotes, ranging from yeast to parasites and humans, the increase in catastrophe rate is driven by upregulation of microtubule-depolymerizing kinesin motors. These motors shorten microtubules, by promoting their catastrophe (Walczak et al., 2013; Welburn, 2013). There are two major families of depolymerizing kinesins: Kinesin-8 and Kinesin-13. Generally, absence of these motors is associated with defects in cell division. Yeast have Kinesin-8 motors only: Kip3 and KlpB from budding yeast and *A.nidulans* respectively, essential for spindle positioning (Cottingham and Hoyt, 1997; Miller et al., 1998; Rischitor et al., 2004; Walczak et al., 1996). Metazoans species such as *D.melanogaster* have both kinesin-8 (Klp67) and kinesin-13 (Klp10, Klp59D) microtubule depolymerases with key roles in mitosis.

In mammalians, the Kinesin-13 family member MCAK, the most potent and best-characterized microtubule-depolymerizing kinesin (Friel and Welburn, 2018), is cytoplasmic localizing to plus ends throughout the cell cycle and to kinetochores in mitosis. *In vitro,* rather than walking on microtubules, MCAK diffuses short distances on the microtubule lattice, although it raises the question of how MCAK reaches the ends of crowded microtubules in cells. This atypical motor also associates with microtubule end-binding (EB) proteins to enhance its localization to microtubule plus ends, utilizing an SxIP-like motif (Honnappa et al., 2009).

Recent work by us and others has shown the Kinesin-8 motor Kif18b shortens astral microtubules in mitosis when it is released into the cytoplasm from the nucleus. It requires both EB proteins and its C-terminal microtubule-binding tail to accumulate at microtubule ends and depolymerize them (McHugh et al., 2018; Stout et al., 2011; Tanenbaum et al., 2011). Absence of Kif18b or MCAK leads to aberrant microtubule growth, defects in spindle assembly and positioning and lagging chromosomes (Domnitz et al., 2012; Huang et al., 2007; McHugh et al., 2018; Rankin and Wordeman, 2010). While Kif18b shortens microtubule length through its depolymerase activity, this activity *in vitro* remains modest with respect to Kinesin-13 motors and does not explain the strong depolymerization associated with Kif18b function *in vivo.* Here using purified proteins, we show that Kif18b transports and accumulates MCAK and EB to microtubule plus ends. We reconstitute an active Kif18b-EB-MCAK network at microtubule plus ends allowing us to dissect the contributions of each factor to the activity of the network. It is only when all three components of the network are present that we observe abolishment of microtubule growth. Overall it is the collective effect of the Kif18b-EB-MCAK network that enables robust change in microtubule dynamics and reduce microtubule length.

## Results

### Kif18 is required for MCAK plus end localization in cells

MCAK and Kif18b both localize to microtubule plus ends and regulate microtubule length in mitosis through a similar pathway (McHugh et al., 2018; Tanenbaum and Medema, 2011). First we defined whether MCAK localization at plus ends depends on Kif18b during mitosis. In HeLa cells, we observed co-localization of MCAK with endogenous EB1 marking the plus ends of astral microtubules (Fig 1a). In cells lacking Kif18b, MCAK was significantly reduced at microtubule plus ends. These data indicate the localization of MCAK to microtubule plus ends was compromised in mitosis when Kif18b is absent in agreement with MCAK plus end reduction after depletion of Kif18b (Tanenbaum et al., 2011).

**Figure 1.**
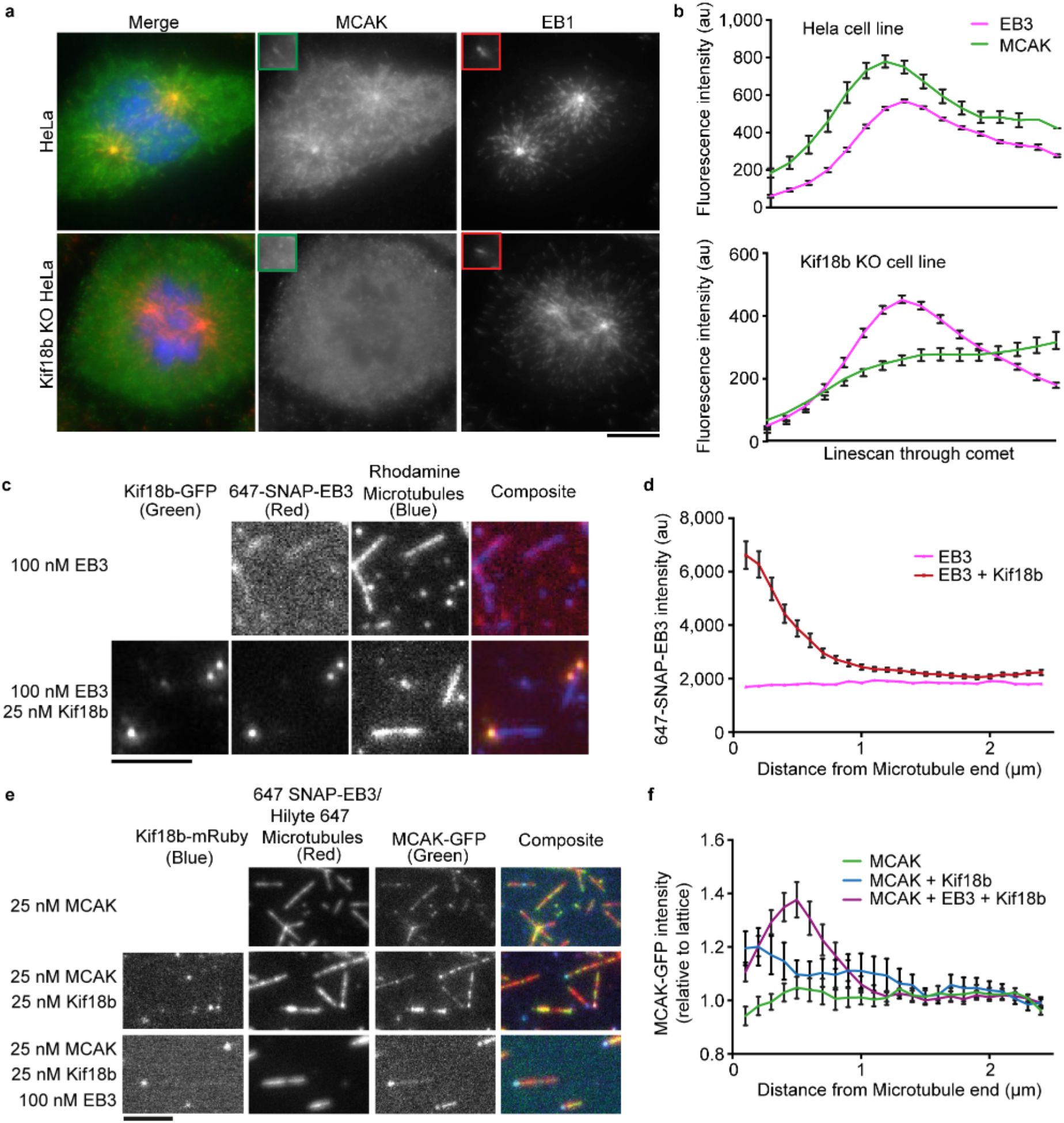
Kif18b increases the localization of MCAK at microtubule plus plus ends. a) Representative images of MCAK (green) and EB3 (red) localization in WT and Kif18b KO Hela cells, b) Quantification of EB3 and MCAK fluorescence intensity at astral microtubule plus ends from 1 experiment (n=30 and 27, respectively), repeated 3 times, c & d) Kif18b increases plus end localization of EB3 on GMPCPP microtubules, (c) Representative images of Kif18b-GFP and Hilyte 647 coupled SNAP-EB3 on GMPCPP stabilized rhodamine labeled microtubules. Scale Bar, 5 μm, (d) Quantification of EB3 fluorescence intensity at the microtubule plus end in the presence (red, n=41) or absence (pink, n=44) of 25 nM Kif18b-GFP (mean and S.D). e & f) Kif18b and EB3 increase the plus end localization of MCAK. (e) Representative images of MCAK-GFP (green) localization on double stable (taxol and GMPCPP) microtubules. Scale Bar 5 μm. (f) Quantification of 25 nM MCAK-GFP fluorescence intensity at microtubule plus ends alone (green, n=76), in the presence of 25 nM Kif18b-mRuby3 (blue, n=90), or 25 nM Kif18b-mRuby3 and 100 nM 647-SNAP-EB3 (purple, n=116).

### Kif18b localizes MCAK and EB3 at stable microtubule plus ends

While MCAK requires Kif18b for plus end localization, the molecular mechanism underlying this dependency is not known. We did not observe any interaction between Kif18b and MCAK in solution despite the weak interaction previously reported (Tanenbaum et al., 2011). We therefore hypothesized Kif18b may interact dynamically with MCAK and EB proteins in the context of the microtubule. We first tested whether Kif18b would interact with EB proteins on the microtubule lattice *in vitro,* using GMPCPP-stabilized microtubules which facilitates our analysis of motor behavior. We generated a GMPCPP–stabilized microtubule lattice, mimicking the GTP cap of microtubules favored by EB proteins. Using *in vitro* reconstitution and multichannel total internal reflection fluorescence (TIRF) microscopy imaging, we imaged microtubules (rhodamine, Hilyte 647), Kif18b (GFP, mRuby3), EB3 (647-SNAP) and MCAK (GFP). While the microtubules in this assay are not polarity-marked, we previously showed Kif18b is a plus end directed motor, accumulating at microtubule plus ends (McHugh et al., 2018). Thus, the Kif18b-GFP accumulation at microtubule tips represents plus end accumulation (Fig 1c, S1a, b). The amount of Kif18b accumulation at microtubule tips was length-dependent (Fig S1a). Recombinant 647-SNAP-EB3 alone uniformly decorated the GMPCPP-stabilized microtubule lattice (Fig 1c, d). Upon addition of 25 nM Kif18b, we observed that 647-SNAP-EB3 accumulated to the plus end of microtubules, co-localizing with Kif18b (Fig 1c, d). Thus, Kif18b is sufficient to bias the localization of EB3 on the microtubule lattice in a nucleotide-independent manner.

Based on the Kif18b-dependent localization of MCAK to microtubule plus ends (Fig 1a, b), we next hypothesized that Kif18b promotes the localization of MCAK to the plus microtubule ends by transporting MCAK along the microtubules. Purified 25 nM MCAK-GFP bound diffusely along the length of double-stabilized (taxol and GMPCPP) microtubules (Fig 1e, f). With the addition of 25 nM Kif18b-mRuby3 the localization of MCAK shifted towards the plus end of the microtubule (Fig 1e, f). The localization of Kif18b-mRuby3 was unchanged by the addition of either MCAK or EB3 (Fig S1b, c). Thus, in our assay, Kif18b promotes the accumulation of both MCAK and EB proteins individually to the plus ends of microtubules. However, MCAK also interacts with EB proteins (Honnappa et al., 2009). In the presence of 100 nM 647-SNAP-EB3 and 25 nM Kif18b-GFP, MCAK plus end accumulation was strongly enhanced (Fig 1e, f).

To test if the Kif18b-dependent localization of MCAK was specific, we incubated the diffusive microtubule crosslinker PRC1-GFP (Subramanian et al., 2010), in the presence of Kif18b. No accumulation of PRC1 to the ends of microtubules was observed, confirming Kif18b-dependent delivery of MCAK and EB3 was specific (Fig S1d). Indeed, the presence of Kif18b at the ends of microtubules led instead to a decrease in PRC1 intensity, perhaps due to competition for space on the lattice in this region. Our data indicate that Kif18b promotes EB and MCAK plus end targeting and that Kif18b and EB proteins function cooperatively to facilitate MCAK microtubule plus end accumulation.

### Kif18b increases MCAK directed movement along the lattice

To test whether Kif18b can directly transport MCAK, we imaged low concentrations of labeled MCAK-GFP on stable microtubules to record single molecule events. MCAK alone diffused on microtubules, with a diffusion coefficient of 5500±500 nm^2^ and remained bound to the lattice for 1.3 seconds on average (Fig 2a, b, Table 1). In the presence of 20 nM Kif18b, MCAK lattice residency increased 1.6-fold and MCAK displayed higher displacement on the lattice (Fig 2a, b, Table 1). At 20 nM Kif18b-mRuby3 and low nanomolar concentrations of MCAK-GFP, required for single molecule imaging, we saw MCAK dwelling at the microtubule ends where the Kif18b concentration will be at its highest (Fig 2c). We also observe rare events of directional movement of MCAK, often at rates of around 300 nm/s corresponding to those seen for single Kif18b motors (McHugh et al., 2018) (Fig 2c). This indicates that the accumulation of MCAK at the plus ends of microtubules in the presence of Kif18b is due to direct and specific transport of MCAK as a cargo of Kif18b. On stable microtubules, Kif18b walks processively and accumulates at microtubule plus ends, using its C-terminal second microtubule-binding domain (McHugh et al., 2018). We found Kif18b continues to gather at microtubule ends whilst also promoting the targeting of MCAK to microtubule plus ends (Fig 1e, S1c). Thus, its C-terminal microtubule-binding domain still likely engages with the microtubule lattice in the presence of MCAK. Overall, the biochemical properties of Kif18b enable the formation of a Kif18b-EB-MCAK network at microtubule plus ends. Multivalent low-affinity interactions between Kif18b, EB and MCAK proteins in the context of the microtubule create an Kif18b-EB-MCAK (KEM) plus end-localized network.

**Table 1.** Residency times and diffusion coefficients fitted to data in Figure 2a and b.

|  | **9.5nM MCAK + 0.5nM MCAK-GFP** |  |
| --- | --- | --- |
|  | **0 nM Kif18b** | **20 nM Kif18b** |
| **Diffusion Coefficient D (nm<sup>2</sup>)** | 5500±500 | 24000±1500 |
| **Residency Time (s)** | 1.3±0.5 | 2.4±0.2 |

**Figure 2.**
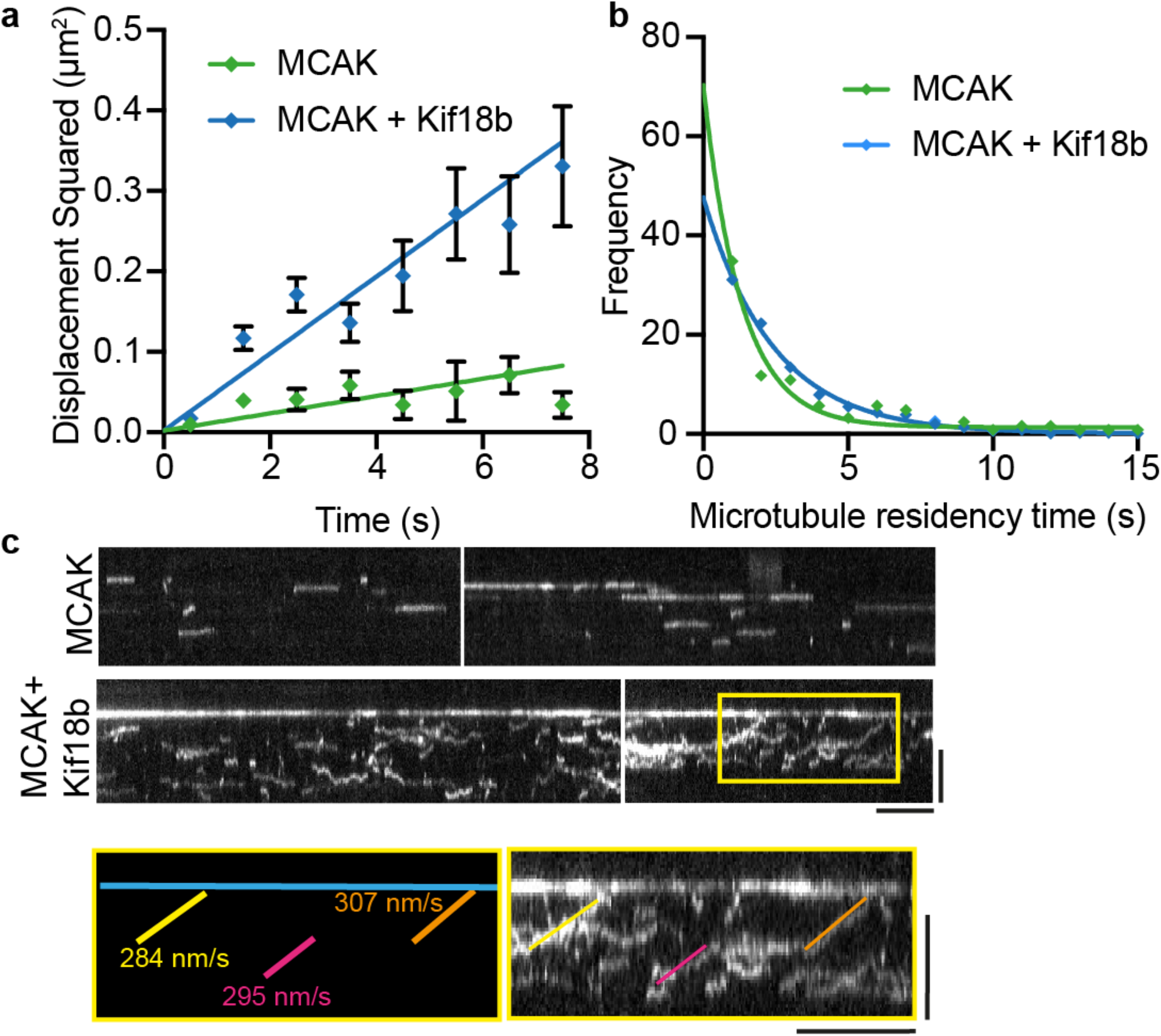
Single MCAK motors show increased lattice diffusion and residency in the presence of Kif18b. a) Mean squared displacement plotted against the time over which it was measured for 0.5 nM MCAK-GFP motors in the presence of 9.5 nM unlabeled MCAK and 0 nM (green, n = 246) or 20 nM Kif18b-mRuby3 (blue, n = 820), mean and S.E. Fitted with a linear curve from which the diffusion coefficient D can be measured (2Dt = <x^2^>, given in Table 1). b) Frequencies of microtubule residency time for MCAK-GFP motors on GMPCPP stable microtubules with 0 nM (green, n=247) and 20 nM (blue, n = 820) Kif18b-mRuby3, mean and S.E. fitted with exponential curves to give mean residency times (see Table 1). c) Example kymographs for 0.5 nM MCAK-GFP and 9.5 nM unlabeled MCAK showing the increase in diffusive behavior in the presence of Kif18b-mRuby3. Scale bars 10 seconds (horizontal) and 3 μm (vertical). Highlighted section shows tracks where MCAK moves directionally towards one end of the microtubule.

### The C terminus of Kif18b inhibits MCAK on stable microtubules

We next tested how these microtubule motors work in the presence of each other to depolymerize stabilized microtubules. Kif18b-GFP does not depolymerize GMPCPP-stabilized microtubules (McHugh et al., 2018). MCAK-GFP alone however efficiently depolymerizes GMPCPP-stabilized microtubules from both ends (Fig 3a) (McHugh et al., 2019). Surprisingly, in the presence of MCAK-GFP and an increasing concentration of Kif18b, the microtubule depolymerization rate was gradually reduced (Fig 2b). The decrease in microtubule end depolymerization was biased and specific to the plus end, where Kif18b-GFP accumulated (Fig 2a). This led to an asymmetry in plus and minus end depolymerization rates, proportional to the concentration of Kif18b (Fig 2c). Overall, Kif18b protects the plus ends of GMPCCP-stabilized microtubules from MCAK-mediated depolymerization. The presence or absence of EB3 which also is known to bind the tail region of Kif18b has no significant effect on the inhibition of MCAK-mediated depolymerization (Fig S2a).

**Figure 3.**
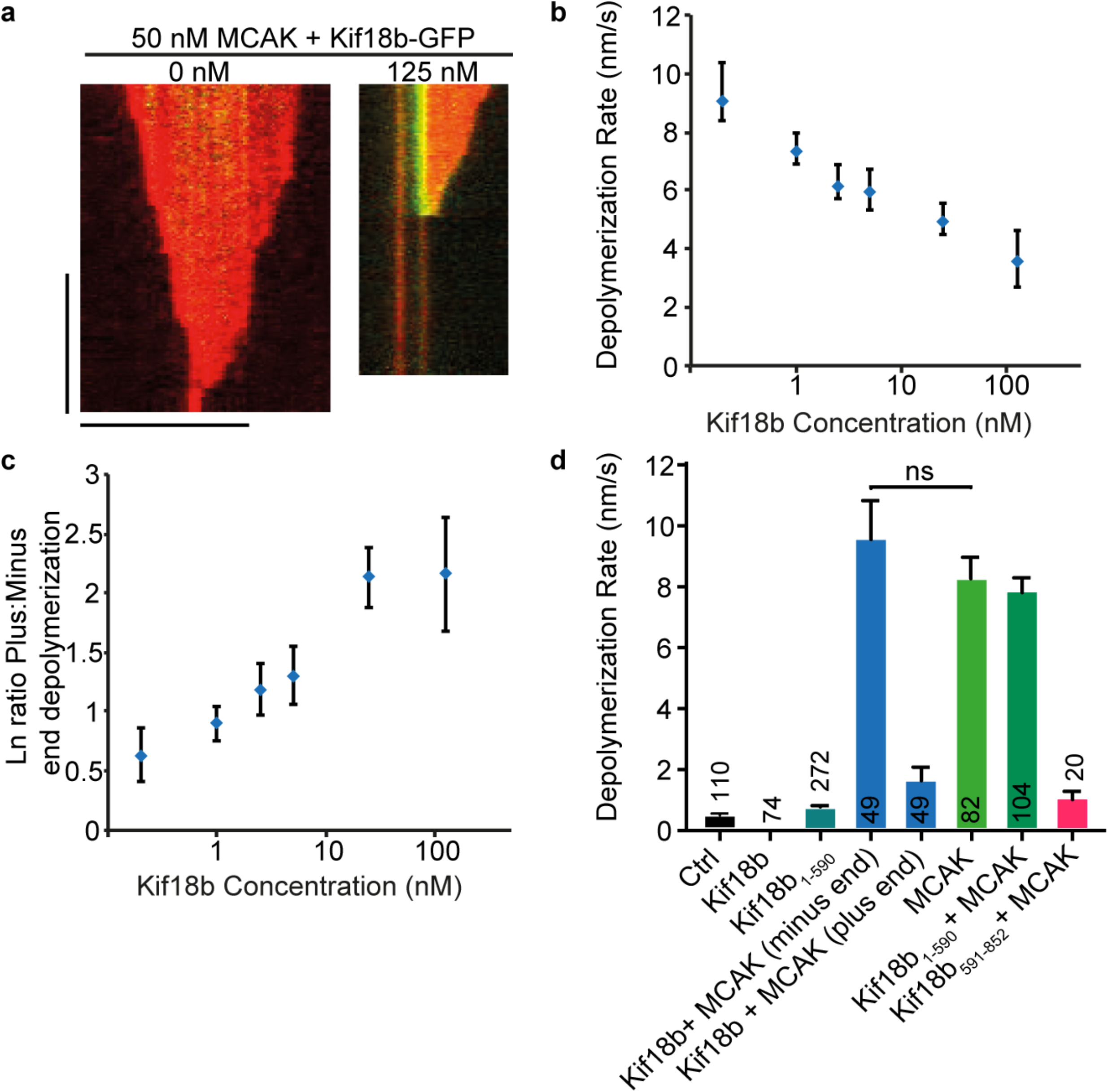
The C-terminal tail of Kif18b stabilizes microtubules to MCAK mediated depolymerization. a) Example kymographs showing the depolymerization of GMPCPP microtubules by 50 nM MCAK-GFP in the presence (right) and absence (left) of 125 nM Kif18b-GFP. Scale bars 1 min (vertical) and 5 μm (horizontal) b) Increasing Kif18b-GFP concentrations lead to lower microtubule depolymerization rates at 50 nM MCAK-GFP. Mean and S.E., n=270. c) The ratio of minus/plus end depolymerization increases with increasing Kif18b concentration. Mean and S.E., n = 270. d) Depolymerization rates for GMPCPP stable microtubules alone, with 50 nM Kif18b-GFP, 50 nM Kif18b_1-590_-GFP, 50nM Kif18b-GFP + 50nM MCAK-GFP (plus and minus end rates), 50 nM MCAK-GFP, 50 nM Kif18b_1-590_-GFP + 50 nM MCAK-GFP and 50 nM MCAK-GFP + 500 nM Kif18b_591-852_. Mean and S.E (see Table S1), n values shown on graph.

We previously showed that the tail of Kif18b, Kif18b_(591-852)_, enables Kif18b to remain bound at the plus ends of microtubules. However it also interferes with the depolymerase activity of the Kif18b catalytic domain (McHugh et al., 2018). We therefore tested whether the C terminus of Kif18b interfered with MCAK-induced microtubule depolymerization. In the presence of 50 nM MCAK and 50 nM Kif18b, the microtubule depolymerization rate of the plus end was severely reduced to 1.61 nm/s while the free minus end depolymerization rate was similar to that with MCAK alone (9.36 nm/s) (Fig 3d, Table S1). A C-terminally truncated Kif18b_590_-GFP was capable of modest microtubule depolymerization, as previously shown (Fig 3d) (McHugh et al., 2018). We found that in the presence of 50nM MCAK and Kif18b_590_-GFP, the microtubule depolymerization rate was similar to the rate in the presence of MCAK alone (Fig 3d). In contrast addition of 500nM Kif18b_(591-852)_ significantly reduced the rate of MCAK-induced depolymerization. It is possible the Kif18b tail (a.a. 591-852) binds to the microtubule lattice to stabilize it, thereby interfering with MCAK-induced-depolymerization. Alternatively, it may bind to MCAK to inhibit it. We could not observe any interaction between MCAK and the Kif18b tail in solution (data not shown). Given that the Kif18b tail binds to microtubules and inhibits the modest depolymerization of the Kif18b motor domains, it is likely to prevent MCAK depolymerization by stabilizing the microtubule lattice. A similar microtubule stabilizing effect has effect has previously been reported for the tail region of the budding yeast kinesin-8, Kip3 (Su et 2011).

Overall, our data indicate that Kif18b enriches MCAK at microtubule plus ends. However, a Kif18b-MCAK complex displays only modest depolymerization activity on stabilized microtubules and greater concentrations of Kif18b inhibit MCAK-mediated microtubule depolymerization further through the action of the Kif18b tail.

In the cellular context, the minus ends of microtubules are protected from depolymerization by minus-end associated proteins such as CAMSAP. Thus, it is unlikely that MCAK can freely depolymerize microtubules primarily from their minus ends. We purified the C terminus of minus-end binding protein CAMPSAP3-C (Roostalu et al., 2018). In the presence of 100 nM CAMSAP3-C, MCAK depolymerization of the microtubule from the minus end was suppressed (Fig S2b). When MCAK, Kif18b and CAMSAP were co-incubated, microtubule depolymerization rates at both ends were reduced. Overall, depolymerization was greater at the plus ends than the minus end of the microtubules, giving a slow net microtubule shrinkage from their plus end termini.

### EB3 recruits MCAK to microtubule ends to increase catastrophe frequency

When Kif18b promotes the targeting of MCAK and EB proteins to the plus ends of stable GMPCPP microtubules, these microtubules are protected from rapid MCAK-mediated depolymerization. However, in cells microtubules are dynamic. Thus, we sought to define the emergent properties of a Kif18b-MCAK-EB network on dynamic microtubules. First, we defined the properties of MCAK on dynamic microtubules both alone and in the presence of EB3. Interestingly, MCAK localized preferentially to growing GDP-microtubule extensions, over the GMPCPP-stabilized seeds that mimic GTP–bound tubulin (Fig 4a). The growth rate of microtubules was unchanged by increasing concentrations of MCAK (Fig S3a). We also found that the growth rates in the presence of MCAK did not vary significantly with microtubule length (Fig S3b). The time for microtubules to undergo catastrophe decreased with increasing MCAK concentration, ranging from 0 to 50 nM (Fig 4b. Table S2). The survival frequency of microtubules not having undergone catastrophe displayed an exponential decay. We were unable to consistently detect microtubule growth events that occurred for a less than 25 seconds so these data were removed from our analysis. The length of microtubules at catastrophe was shorter in the presence of MCAK with a decay constant of 1170 nm for 50 nM MCAK against 2510 nm for the control (Figure 4c, Table S2). This is in agreement with a previous study (Gardner et.al. 2011).

**Figure 4.**
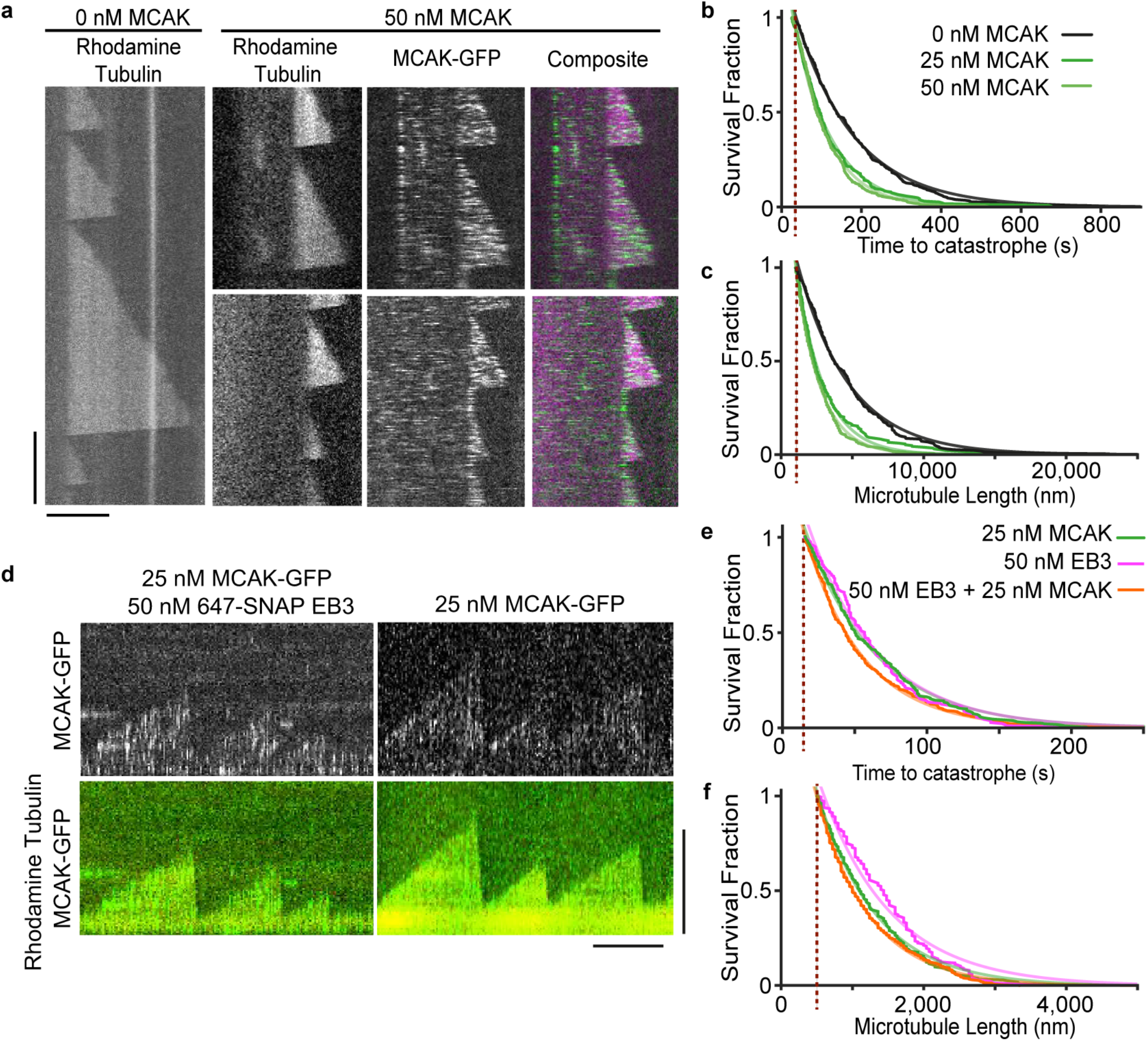
MCAK and EB3 increase microtubule catastrophe frequency and decrease microtubule length. a) Representative kymographs of dynamic microtubules in the presence of 50 nM MCAK-GFP. Scale 2 mins (vertical) and 10 μm (horizontal) b, c) MCAK decreases catastrophe time and length. Survival fraction of microtubule extensions until catastrophe time (b) and catastrophe length (c) with increasing MCAK-GFP concentration. 0 nM (black), 25 nM (green), 50 nM (light green). Curves are fit by exponential decays, n > 220 for each condition, parameters are given in Table S2. Red dotted line indicates time resolution cut of at 25 seconds (b) and length resolution cut off at 1000 nm (c). d) Representative kymographs of dynamic microtubules in the presence of 25 nM MCAK-GFP and 50 nM 647-SNAP-EB3. Scale 10 μm (vertical) and 2 mins (horizontal). e) Addition of EB3 to MCAK has little effect on catastrophe time and length. Survival fraction of microtubule extensions until catastrophe time (e) and length (f) with 25 nM MCAK (green), 50 nM EB3 (pink) or 25 nM MCAK and 50 nM EB3 (orange). Curves are fit by exponential decays, n > 175 for all conditions, parameters are given in Table S2. Red dotted line indicates time resolution cut of at 25 seconds (e) and length resolution cut of at 500 nm (f).

We next analyzed whether microtubule dynamics in the presence of MCAK were altered by the addition of EB3 (Fig 4d). When compared with MCAK alone, microtubule growth rate was unchanged in the presence of 50 nM EB3, while catastrophe time and microtubule length were shortened (Fig 4d-f, S3c). Additionally, we could observe enrichment of MCAK to the growing ends of microtubules in the presence of EB3 (Fig 4d, S3d). In total these data indicate EB3 promotes MCAK plus end targeting to increase microtubule catastrophe and shorten microtubules in broad agreement with previous work by Montenegro et al. 2010.

### The collective behavior of Kif18b and MCAK causes increase in microtubule catstrophe rate

We next tested whether Kif18b and MCAK influence microtubule dynamics synergistically or additively. Using 25 nM of each motor, we compared the effect of the motors alone and in combination. At 25 nM, Kif18b has a modest effect, increasing catastrophe frequency and growth rate leading to a slightly reduced microtubule extension length (McHugh et.al. 2018). MCAK decreased the time to catastrophe for growing microtubules (Fig 5b) and this occurred to a greater extent than for Kif18b (Fig 5a, b). However, in the presence of both Kif18b and MCAK, time to catastrophe decreased 1.8-fold from 44.9 seconds for MCAK alone to 25.5 seconds when in combination with Kif18b (Fig 5b, Table S4). This decrease in growing time was the result of synergistic activity of MCAK and Kif18b. Growth rate was unchanged in the presence of the distinct motors. Microtubule length was strongly reduced in the presence of MCAK and further reduced when both Kif18b and MCAK were present (Figure 5c). We observed the localization of MCAK-GFP on dynamic microtubules and found that similarly to stable microtubules, the presence of Kif18b enhanced the amount of MCAK at the ends of growing microtubules (Fig 5d, e). This was specific to microtubule plus ends with no change to MCAK-GFP residency seen on the lattice. Overall, these data indicate that Kif18b and MCAK cooperate to increase catastrophe rate and thus reduce microtubule length.

**Figure 5.**
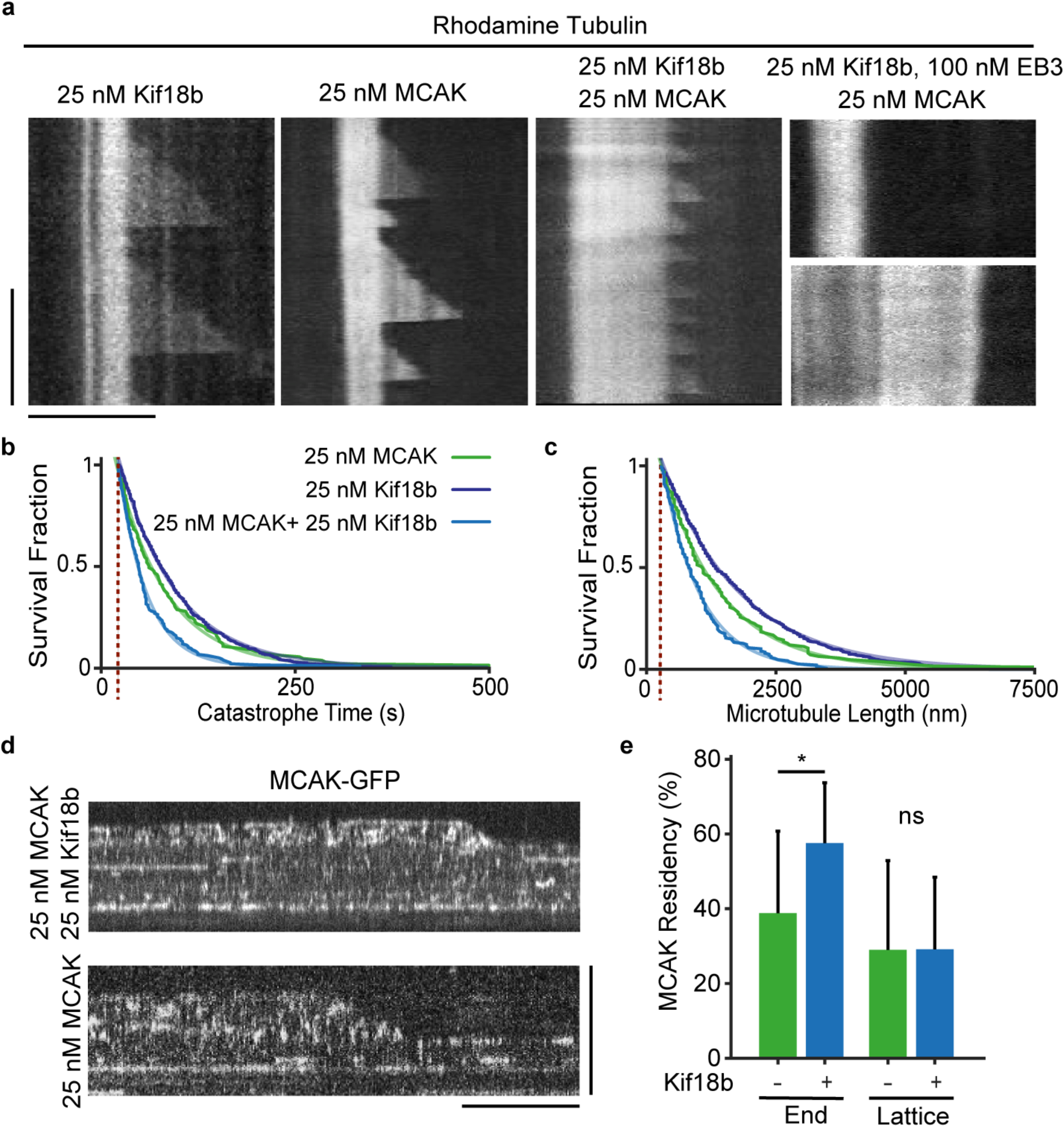
Cooperation of Kif18b, MCAK and EB3 lead to high catastrophe frequencies and inhibit microtubule dynamics. a) Representative kymographs of dynamic microtubules in the presence of 25 nM MCAK-GFP, 25 nM Kif18b-mRuby3 and 100 nM 746-SNAP-EB3. Scale, 5 mins (vertical) and 10 μm (horizontal) b, c) MCAK and Kif18b together decrease both catastrophe time and length. Survival fraction of microtubule extensions until catastrophe time (b) and catastrophe length (c) with 25 nM MCAK (green), 25 nM Kif18b (dark blue) and 25 nM Kif18b + 25 nM MCAK (blue). Curves are fit by exponential decays, n > 110 for all conditions, parameters are given in Table S4. Red dotted line indicates time resolution cut off at 25 seconds (b) and length resolution cut off at 500 nm (c). d) Representative kymograph showing 25nM MCAK-GFP localization on dynamic microtubules in the presence of 25nM Kif18b-mRuby3. Scale 10 seconds (vertical) and 10 μm (horizontal) e) Percentage of time that MCAK-GFP is resident at the dynamic microtubule plus end, within 160nm, or on the lattice further than 1 μm from plus end for 25nM MCAK alone (green, n=17) or with 25 nM Kif18b (blue, n=11), mean and S.D.

### Kif18b, EB3 and MCAK work collectively to inhibit microtubule growth

We next defined the emergent properties of the Kif18b-EB-MCAK (KEM) network with respect to microtubule dynamics. Only in the presence of all three proteins we observed no microtubule growth (Fig 5a). Under this condition specifically, we also noted that some microtubules displayed slow depolymerization of the GMPCPP-stabilized seed (Fig 5a). When EB3 and MCAK or MCAK and Kif18b were combined, growth rate was largely unchanged with catastrophe rate (Table S2, S3). EB3 alone had only a mild effect above that of MCAK on microtubule dynamics (Fig 4d-f), while Kif18b and MCAK together had a modest effect. The addition of a third component to the assay containing either MCAK and EB3 or MCAK and Kif18b, dramatically changes the microtubule dynamics to completely prevent any microtubule growth. Together, the KEM network work in an integrated manner to robustly depolymerize microtubules.

## Discussion

Microtubule dynamics dramatically increase at mitotic onset, to remodel the microtubule cytoskeleton and assemble a bipolar spindle. In the absence of Kif18b or MCAK, mitotic microtubules remain long, leading to spindle assembly and positioning defects. Using single molecule imaging, we demonstrated here that Kif18b interacts with, and transports MCAK on the microtubule lattice to microtubule plus ends *in vitro.* Kif18b also brings EB3 to microtubule plus ends in a nucleotide-independent manner. These three factors collectively form a network of multivalent weak interactions with potent microtubule shortening activity at microtubule plus ends, which is necessary for correct formation and positioning of the spindle. Our data challenge previous models that postulate MCAK diffuses on microtubules to reach microtubule ends (Cooper et al., 2010; Helenius et al., 2006). While 1-dimensional diffusion on the microtubule lattice is an efficient way to reach a microtubule end for short distances, diffusion is not necessarily an efficient way to reach specifically the plus end of a microtubule in a cell. The diffusion rate of MCAK was calculated as 5,500 nm^2^/s with an off-rate of 0.77 per second (Table 1). This would mean that a single motor would on average explore just 84 nm of the microtubule lattice with each encounter. In the presence of Kif18b, this is increased 4-fold, with MCAK diffusing an average of 240 nm, thereby increasing the probability MCAK would reach or get close to a microtubule plus end. Diffusion alone does not explain how MCAK localizes primarily to microtubule plus ends on crowded microtubules in cells. In addition, we found that *in vitro* MCAK prefers to bind the GDP over the GTP lattice, again suggesting that in cells additional factors are required to target MCAK to the ends of growing microtubules which are necessarily in the GTP bound state (Fig 6a, b).

**Figure 6.**
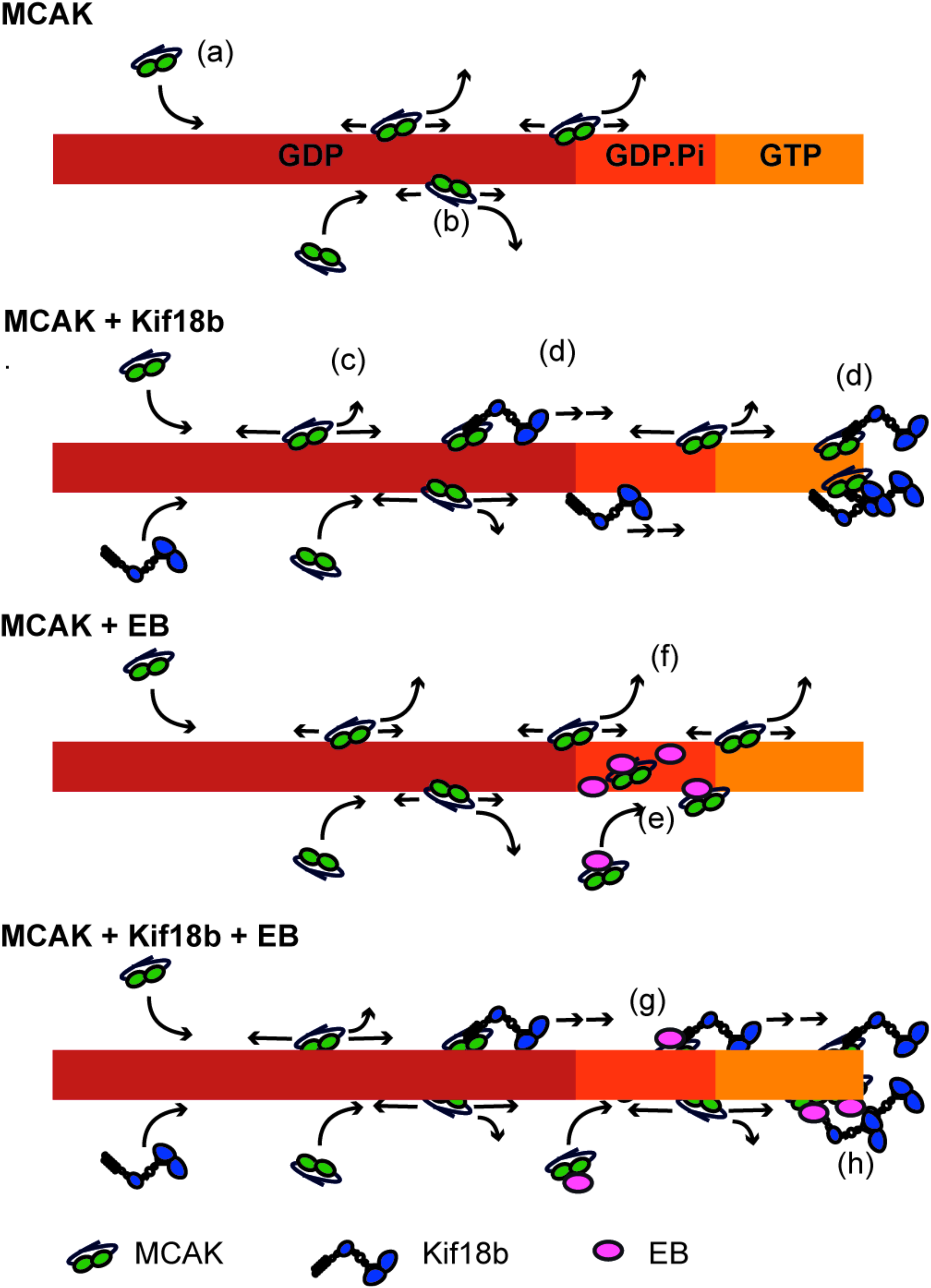
Model of the Kif18b-EB-MCAK network at dynamic microtubule ends. MCAK alone binds predominantly to the GDP microtubule lattice (a) and diffuses for short distances before detaching (b). Few MCAK motors reach the microtubule end. Upon addition of Kif18b, MCAK diffuses further on the microtubule lattice (c), rare transport events also occur (d). This leads to more MCAK arriving at the microtubule ends. MCAK binds to EB proteins in solution and on the microtubule lattice where EBs recognize the nucleotide state of the microtubule near the end (e). This increases the amount of MCAK diffusing near the end of the microtubule. When both EB and Kif18b proteins are present, Kif18b can bind to both EB and MCAK (f) this strengthens the weak interaction between MCAK and Kif18b and allows Kif18b to transport both EB and MCAK to the very end of the microtubule (g).

While Kif18b alone is a weak microtubule depolymerase, our results indicate it is a major microtubule depolymerization factor in mitosis once released from the nucleus. It delivers MCAK and EB proteins to microtubule ends, where they work together in an integrated manner to prevent microtubule growth. We propose that this cooperative mechanism enhances MCAK concentration at microtubule plus ends in three ways: firstly, through increasing the amount of time that MCAK spends bound to the microtubule (Fig 6c); secondly, through the direct targeting of MCAK to the microtubule plus end either by transport (Kif18b; Fig 6d) or through binding in the tip region (EB proteins; Fig 6e); thirdly, through an increase in the multivalent interactions of MCAK with EB3, Kif18b and the microtubule (Fig 6f). In the context of the KEM network, MCAK and Kif18b may also gain additional functions, as our study suggest. Overall, this KEM network displays strong depolymerization activity that cannot be imparted to the sum of the individual components, highlighting the emergent properties of the network.

Our data suggest Kif18b plays a critical role as an integration platform to modulate the depolymerase activity of the network. Indeed, the Kif18b C-terminal tail binds microtubules, EB3 and MCAK allowing it to integrate multiple signals and to modulate the output of the complex’s activity. We previously showed the Kif18b tail is essential for Kif18b plus end targeting and has a complex regulatory role on microtubule dynamics with opposing effects of Kif18b on dynamic versus stable microtubules (McHugh et al., 2018; Tanenbaum and Medema, 2011). Kif18b shows length dependent accumulation at microtubule plus ends *in vitro* through its C terminus (Fig S1a), a hallmark of Kinesin-8 motors (Shrestha et al., 2018), which would enable the network to preferentially target and depolymerize longer microtubules. Additionally, two distinct populations of Kif18b decorating astral microtubule plus ends have been reported; one population of Kif18b interacts and colocalizes with EB1, while the other precedes EB1 at the plus end of the microtubule (Shin et al., 2015). Kif18b may recognize end structures in both an EB1-dependent and - independent fashion. In cells EB proteins do not bind to the terminal tubulin dimers but are instead displaced ~100 nm from the plus end (Nakamura et. al. 2012). It is possible that on dynamic microtubules Kif18b helps localize EB independently of nucleotide state and therefore brings the MCAK-EB network closer to the terminal tubulin dimers to promote depolymerization. This would ensure that all three proteins are present together to promote robust microtubule depolymerization. Importantly, each interaction within this KEM network is individually regulated by mitotic phosphorylation, a rapid and reversible post-translational modification, while Kif18b is confined to the nucleus before mitosis and irreversibly degraded at the end of mitosis (Lee et al., 2010; McHugh et al., 2018; Tanenbaum et al., 2011). Thus, the integrity and activity of this depolymerizing network is tightly regulated to allow for spatial, local and temporal regulation of microtubule dynamics during mitosis.

Finally our work highlights the importance of studying motors not only in isolation but examining their collective behavior and the underlying emerging function. How the cargos influence motor activity remains a poorly understood phenomenon. Rather than having a microtubule motor that provides transport and passive cargos, we reveal here how the three components of mitotic-specific KEM network, two of which are motors, act in an integrated manner to provide a novel depolymerizing function with temporal and spatial layers of control.

## Author contributions

JW conceived the study. TM and JW acquired, analyzed and interpreted the data. TM and JW wrote the manuscript.

## Competing interests

The authors have no competing interests to declare.

## Acknowledgements

J. W. is supported by a Wellcome Senior Reseach Fellowship (207430). We thank Agata Gluszek for the EB3-SNAP-His plasmid and Dave Kelly for COIL support. The Wellcome Trust Centre for Cell Biology is supported by core funding from the Wellcome Trust (203149).

## Methods

### Cloning

The Sf9 insect cell expression constructs for producing full-length human Kif18b-GFP, Kif18b_1-590_-GFP and MCAK-GFP with a C-terminal 6xHis have been described elsewhere (McHugh et al., 2018; Talapatra et al., 2015). For Kif18b-mRuby3-His, Kif18b was cloned in to pFl-mRuby3-His using restriction enzymes. mRuby3 was synthesized after codon optimization for expression in insect cells. Kif18b_591-852_ was cloned into a pET 3atr with an N-terminal 6xHis. EB3 was cloned into vectors containing a SNAP-His or GFP-His tag. The His-TEV-GFP-PRC1 and StrepTagII-mGFP-G5A-CAMSAP3-C (pCT011) plasmids were gifts from T. Kapoor and T. Surrey and are described in (Roostalu et al., 2018; Subramanian et al., 2010), respectively.

### Cell culture

HeLa cells were used and maintained in DMEM (Lonza) supplemented at 37°C in a humidified atmosphere with 5% CO_2_. The generation of the Kif18b knockout cell line is described in (McHugh et al., 2018). Cells are monthly checked for mycoplasma contamination (MycoAlert detection kit, Lonza).

### Immunofluorescence and microscopy

Cells were washed in PBS and fixed in 3.8% formaldehyde in PHEM buffer (60 nM Pipes, 25 mM HEPES, 10 mM EGTA, 2 mM MgSO_4_, pH 7.0) for 10 minutes. Immunofluorescence in human cells was conducted using antibodies against mouse EB1 (BD transduction laboratories, 1:400) and a custom rabbit MCAK antibody (1:500-1000). The MCAK antibody, raised against GFP-MCAK, was generated by Eurogenetec using the Speedy program. Over time, the antibody became rapidly unstable and non-specific. Images were acquired using a DeltaVision core microscope (Applied Precision) equipped with a CoolSnap HQ2 CCD camera. 10-20 z-sections were acquired at 0.2-0.5 μm. Comet intensities were quantified as previously described (McHugh et al., 2018). Experiments were repeated three times.

### Protein expression and purification

Kif18b, MCAK and CAMSAP proteins were expressed using a baculovirus expression system, Sf9 cells were infected with virus for each construct for 60-72 hours. EB3 and PRC1 proteins were expressed in BL21 Codon+ E.coli cells overnight at 18°C. CAMSAP proteins were purified as described in Roostalu et al., 2018. His-tagged Kif18b and MCAK proteins were purified as previously described (Talapatra et al., 2015). His-tagged GFP-EB3, SNAP-EB3 and GFP-PRC1 proteins were purified using a HisTrap HP column and a gradient elution. His-SNAP-EB3 proteins were then coupled to SNAP-Cell 647-SiR dye (NEB) by incubation at 37°C for 30 minutes. All proteins were further purified using gel filtration chromatography pre-equilibrated in gel filtration buffer (25 mM Pipes, pH 6.9, 150 mM NaCl, 300 mM KCl, 5 mM β-mercaptoethanol, 1 mM MgCl_2_, 1 mM Na-EGTA, and 1 mM ATP). Porcine brain tubulin was purified as described (Castoldi and Popov, 2003).

#### Sample preparation for TIRF microscopy

Samples were prepared as detailed in (McHugh et al., 2018). An ATP concentration of 2 mM and a temperature of 30°C were used for all experiments. The stated concentrations of protein (or an equivalent volume of gel filtration buffer) were added to the final assay buffer (BRB80 with 1 mM ATP, 0.5 mg/ml casein, and an oxygen-scavenging system [0.2 mg/ml glucose oxidase, 0.035 mg/ml catalase, 4.5 mg/ml glucose, and 140 mM *β* - mercaptoethanol]), slides were sealed with Valap and imaged immediately. For dynamic microtubule experiments, the assay buffer also contained 7-12 μM tubulin (6% rhodamine-label), 1mg/ml casein and 1mM GTP. Experiments which are compared (in the same graph or table) in this paper were done in parallel using the same ‘master mix’ assay buffer with at least three independent repeats, except for Fig 5a, Kif18b, EB3 + MCAK on dynamic microtubules which was repeated twice and Fig S2a which has one repeat.

#### TIRF Microscopy Imaging

Imaging was performed on a Zeiss Axio Observer Z1 TIRF microscope using a 100 × NA 1.46 objective and a Photometrics Evolve Delta electron-multiplying charge-coupled device camera controlled by Zeiss Zen Blue software. For single molecule experiments a 1.6x tube lens was used, an Optosplit III beam-splitter in bypass mode inserted before the camera provided a further 2x magnification. Depolymerization assays were performed over 10 minutes at 1 fps or 0.5 fps for two colour imaging, this had no effect depolymerization rates. Microtubule dynamics assays were performed over 15 minutes in either one or two colours at 0.3 fps, MCAK single molecule and plus end residency assays were performed in the GFP Channel only at 10 fps.

#### Image analysis

Kymographs were produced using ImageJ (National Institutes of Health), MCAK single molecule behaviour and microtubule growth rates, catastrophe length and catastrophe time were manually measured from these kymographs. Line scan intensity measurements along microtubules were taken using a fixed threshold in the microtubule fluorescence channel (Rhodamine or Hilyte 647) to specify the microtubule ends. Microtubule end residency was calculated from kymographs using line scans, at the microtubule end or on the lattice 1200 nm from the plus end, with a line width of 160 nm. The residency was defined as the percentage of time points where the intensity of the line was above the maximum of the background measured for each kymograph. Microtubule dynamics data were analysed using MATLAB R2020a to fit exponential decay curves and normal distributions to catastrophe frequency and growth rate data respectively. Graphpad Prism 8 was used for all other data analysis.

#### Statistics and reproducibility

Statistical analyses were performed using Prism 6.0 (GraphPad Software) or MATLAB R2020a software. No statistical method was used to predetermine sample size. No samples were excluded from the analyses. The investigators were not blinded to allocation during experiments and outcome assessment. All experiments were performed and quantified from at least three independent experiments (unless specified otherwise), and representative data are shown.

## Figure legends

**Supplementary Figure 1**

a) Dependence of maximum fluorescence intensity of Kif18b-GFP at microtubule plus ends on the length of microtubule, 0.5 μm bins, mean and S.E. n=42. b) Quantification of the GFP fluorescence intensity along the microtubule in the presence of 100 nM 647-SNAP-EB3 and 25 nM (red, n = 41) or 0 nM (pink, n = 44) of Kif18b-GFP (mean and S.D). c) Quantification of mRuby3 fluorescence intensity along the microtubule in the presence of 25 nM Kif18b-mRuby3 alone (dark blue, n = 64), with 25 nM MCAK-GFP (blue, n = 90), with 25 nM MCAK-GFP and 100 nM 647-SNAP-EB3 (purple, n = 116). d) Quantification of GFP fluorescence intensity along the microtubule in the presence of 100 nM GFP-PRC1 and 25 nM (dark blue, n = 26) or0 nM (green, n = 26) of Kif18b-mRuby3 (mean and S.D).

**Supplementary Figure 2**

a) Depolymerisation rates of GMPCPP stable microtubules by 200nM MCAK alone (n=70) with 25nM Kif18b (n=56) and with 200nM EB3 and 25nM Kif18b (n=52). Mean and S.E., data from one repeat only. b) Depolymerization rates of GMPCPP stable microtubule plus and minus plus ends by 10 nM MCAK-GFP in the presence of 100 nM mCherry-CAMSAP (n = 29), 10 nM Kif18b-GFP (n = 36) or 100 nM mCherry-CAMSAP + 10 nM Kif18b-GFP (n = 35). Mean and S.E.

**Supplementary Figure 3**

a) Growth rate of microtubules is not affected by MCAK-GFP concentration, 0 nM (black, n = 272), 25 nM (green, n = 244), 50 nM (light green, n = 404). Gaussian fitted to 5 nm/s bins, parameters given in Table S2. b) Growth rates remain constant with microtubule length for MCAK, mean and S.D., bin widths are 1500 nm, dotted lines show linear fits to the binned data. c) Growth rate of microtubules with 25 nM MCAK-GFP is increased upon addition of 50 nM EB3. Gaussian fitted to 5 nm/s bins, parameters given in Table S3, 25 nM MCAK (green, n = 313), 50 nM EB3 (pink, n = 183), 25 nM MCAK and 50 nM EB3 (orange, n = 398). d) Percentage of time that MCAK-GFP is resident at the dynamic microtubule plus end, within 160nm, or on the lattice greater than 1 μm from plus end for 25nM MCAK alone (green, n = 9) or with 50 nM EB3 (pink, n = 8), mean and S.D.

**Supplementary Figure 4**

a) Growth rate of microtubules are not affected by MCAK-GFP or Kif18b-mRuby3 concentration, 25 nM MCAK (green, n = 130), 25 nM Kif18b (dark blue, n = 615) and 25 nM Kif18b + 25 nM MCAK (blue, n = 192). Gaussian fitted to 5 nm/s bins, parameters give in Table S4.

**Supplementary Table 1.** Summary of depolymerization rates (mean and S.E.) and n values for the data shown in Figure 3d.

|  | **Depolymerization Rate (nm/s)** | **N** |
| --- | --- | --- |
| **Ctrl** | 0.411±0.041 | 110 |
| **Kif18b<sub>FL</sub>** | 0.007±0.001 | 74 |
| **Kif18b<sub>1-590</sub>** | 0.716±0.045 | 272 |
| **Kif18b<sub>FL</sub> + MCAK (+ end)** | 1.61±0.435 | 49 |
| **Kif18b<sub>FL</sub> + MCAK (- end)** | 9.36±1.25 | 49 |
| **MCAK** | 8.11±0.68 | 82 |
| **Kif18b<sub>1-590</sub> + MCAK** | 7.70±0.47 | 104 |
| **Kif18b<sub>591-852</sub> + MCAK** | 1.04±0.26 | 20 |

**Supplementary Table 2.**
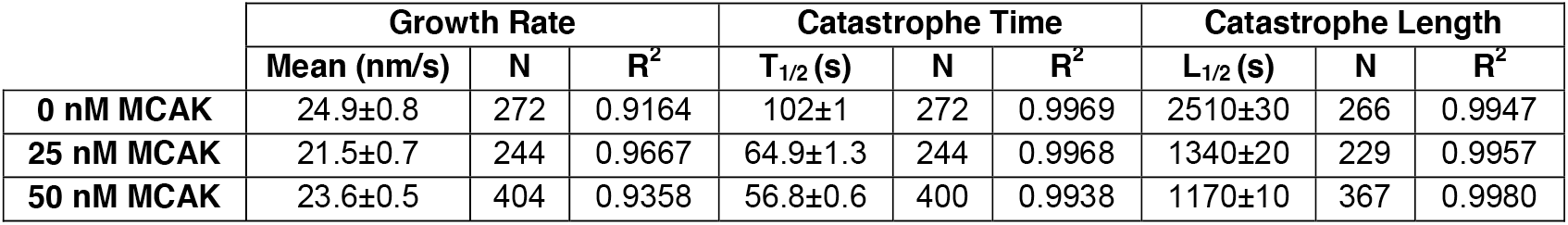
Microtubule dynamics parameters in the presence of MCAK. A tubulin concentration of 12 μM was used. Parameters for data in Figure 4b and c and Supplementary Figure 3a. Growth rates were fitted by a normal distribution giving mean and S.E. Catastrophe Time and Catastrophe Length are fit by exponentials, half-lives of fitted curves with 95% C.I. are given. N values and R-squared values for data and fitted curves are also given.

|  | **Growth Rate** |  |  | **Catastrophe Time** |  |  | **Catastrophe Length** |  |  |
| --- | --- | --- | --- | --- | --- | --- | --- | --- | --- |
|  | **Mean (nm/s)** | **N** | **R<sup>2</sup>** | **T<sub>1/2</sub> (s)** | **N** | **R<sup>2</sup>** | **L<sub>1/2</sub> (s)** | **N** | **R<sup>2</sup>** |
| **0 nM MCAK** | 24.9±0.8 | 272 | 0.9164 | 102±1 | 272 | 0.9969 | 2510±30 | 266 | 0.9947 |
| **25 nM MCAK** | 21.5±0.7 | 244 | 0.9667 | 64.9±1.3 | 244 | 0.9968 | 1340±20 | 229 | 0.9957 |
| **50 nM MCAK** | 23.6±0.5 | 404 | 0.9358 | 56.8±0.6 | 400 | 0.9938 | 1170±10 | 367 | 0.9980 |

**Supplementary Table 3.** Microtubule dynamics parameters in the presence of MCAK and EB3. Parameters for data in Figure 4e and f and Supplementary Figure 3c. Growth rates were fitted by a normal distribution giving mean and S.E. Catastrophe Time and Catastrophe Length are fit by exponentials, half-lives of fitted curves with 95% C.I. are given. N values and R-squared values for data and fitted curves are also given.

|  | **Growth Rate** |  |  | **Catastrophe Time** |  |  | **Catastrophe Length** |  |  |
| --- | --- | --- | --- | --- | --- | --- | --- | --- | --- |
|  | **Mean (nm/s)** | **N** | **R<sup>2</sup>** | **T<sub>1/2</sub> (s)** | **N** | **R<sup>2</sup>** | **L<sub>1/2</sub> (s)** | **N** | **R<sup>2</sup>** |
| **25 nM MCAK** | 15.7±0.3 | 313 | 0.9657 | 33.7±0.3 | 283 | 0.9966 | 575±5 | 247 | 0.9968 |
| **50 nM EB3** | 21.8±0.4 | 183 | 0.9777 | 32.0±0.8 | 178 | 0.9831 | 705±20 | 181 | 0.9764 |
| **25 nM MCAK + 50 nM EB3** | 19.0±0.2 | 398 | 0.9704 | 26.6±0.2 | 343 | 0.9982 | 520±5 | 330 | 0.9983 |

**Supplementary Table 4.** Microtubule dynamics parameters in the presence of MCAK and Kif18b. Parameters for data in Figure 5b and c and Supplementary Figure 4a. Growth rates were fitted by a normal distribution giving mean and S.E. Catastrophe Time and Catastrophe Length are fit by exponentials, half-lives of fitted curves with 95% C.I. are given. N values and R-squared values for data and fitted curves are also given.

|  | **Growth Rate** |  |  | **Catastrophe Time** |  |  | **Catastrophe Length** |  |  |
| --- | --- | --- | --- | --- | --- | --- | --- | --- | --- |
|  | **Mean (nm/s)** | **N** | **R<sup>2</sup>** | **T<sub>1/2</sub> (s)** | **N** | **R<sup>2</sup>** | **L<sub>1/2</sub> (s)** | **N** | **R<sup>2</sup>** |
| **25 nM MCAK** | 17.6±0.3 | 130 | 0.9154 | 44.9±0.8 | 116 | 0.9959 | 809±10 | 124 | 0.9979 |
| **25 nM Kif18b** | 17.6±0.2 | 615 | 0.9110 | 51.0±0.2 | 603 | 0.9988 | 1050±1 | 608 | 0.9986 |
| **25 nM MCAK + 25 nM Kif18b** | 18.0±0.3 | 192 | 0.9186 | 25.5±0.3 | 173 | 0.9958 | 510±6 | 167 | 0.9986 |

## Notes

### Competing Interest Statement

The authors have declared no competing interest.

